# Clec12a tempers inflammation while restricting expansion of a colitogenic commensal

**DOI:** 10.1101/2023.03.16.532997

**Authors:** Tyson R. Chiaro, Kaylyn M. Bauer, Kyla S. Ost, Emmanuel Stephen-Victor, Morgan C. Nelson, Jennifer H. Hill, Rickesha Bell, Morgan Harwood, Warren Voth, Taylor Jackson, Kendra A Klag, Ryan M. O’Connell, W. Zac Stephens, June L. Round

**Affiliations:** University of Utah School of Medicine, Department of Pathology, Division of Microbiology and Immunology, UT 84211

## Abstract

Regulation of the microbiota is critical to intestinal health yet the mechanisms employed by innate immunity remain unclear. Here we show that mice deficient in the C-Type-lectin receptor, Clec12a developed severe colitis, which was dependent on the microbiota. Fecal-microbiota-transplantation (FMT) studies into germfree mice revealed a colitogenic microbiota formed within Clec12a^-/-^ mice that was marked by expansion of the gram-positive organism, *Faecalibaculum rodentium*. Treatment with *F. rodentium* was sufficient to worsen colitis in wild-type mice. Macrophages within the gut express the highest levels of Clec12a. Cytokine and sequencing analysis in Clec12a^-/-^ macrophages revealed heighten inflammation but marked reduction in genes associated with phagocytosis. Indeed, Clec12a^-/-^ macrophages are impaired in their ability to uptake *F. rodentium.* Purified Clec12a had higher binding to gram-positive organisms such as *F. rodentium*. Thus, our data identifies Clec12a as an innate immune surveillance mechanism to control expansion of potentially harmful commensals without overt inflammation.

## INTRODUCTION

The composition and function of the microbiota can have a profound impact on mammalian health and disease (Fassarella et al., 2021). This has been well documented for inflammatory bowel disease (IBD), where most mouse models of IBD are sensitive to changes in microbiota composition (Gkouskou et al., 2014). Understanding the factors that influence and instruct microbiota stability will be important as microbiota-based therapies are developed (Sharma et al., 2020; Sorbara and Pamer, 2022). The microbiota is influenced by multiple parameters including diet, medications, and the immune system. In recent years, adaptive immune components, especially IgA, have been shown to influence microbiota composition (Nakajima et al., 2018; Okai et al., 2016; Yang and Palm, 2020). However, less is known about how specific innate immune receptors influence homeostatic microbiota composition.

Genetic polymorphisms in genes regulating microbial recognition and mucosal immunity are significant risk alleles for IBD, including the autophagy protein ATG16L1 (Cadwell et al., 2008; Liu et al., 2015; Rioux et al., 2007; Wellcome Trust Case Control, 2007). In a recent study exploring myeloid gene expression changes in individuals with the ATG16L1T300A polymorphism, the inhibitory C-type lectin receptor (CLR), Clec12a, was identified to be significantly suppressed in these cells (Begun et al., 2015). This same study demonstrated that Clec12a functions in defense of pathogenic infection through association with the autophagy pathway. These data suggest that Clec12a downregulation in individuals with IBD might influence disease severity and/or development. Despite this evidence, little is known about the immune regulation mechanisms exerted by Clec12a during colitis.

C-type lectins are a diverse group of soluble or membrane bound molecules capable of binding both host-or microbial-derived, lipids, proteins and carbohydrates (Brown et al., 2018). Mincle (*Clec4e*) and Dectin-1 (*Clec7a*) are two CLRs that influence specific members of the microbiota to maintain intestinal homeostasis (Iliev et al., 2012; Martinez-Lopez et al., 2019). However, with over 1,000 different C-type lectins, much remains to be discovered within this protein family. Clec12a belongs to the natural killer receptor family and is one of only two C-type lectins to contain an immune-tyrosine inhibitory motif (ITIM) capable of restricting inflammatory signaling pathways (Brown *et al*., 2018; Marshall et al., 2004). Clec12a recognizes uric acid which is a signal of cellular damage(Neumann et al., 2014). Clec12a is most highly expressed on innate immune cells, and in neutrophils it acts to reduce activation and downregulate inflammatory cytokine production upon uric acid recognition (Neumann *et al*., 2014). Conversely, Clec12a promotes inflammation by permitting mono-nuclear phagocytes (MNPs) transmigration across the blood-brain barrier during an autoimmune model of multiple sclerosis and potentiating type I interferon signaling in response to viral infection (Li et al., 2019; Sagar et al., 2017). Clec12a is also capable of recognizing the plasmodial product, hemozoin, which has been shown to increase CD8^+^ T cells in the brain following *Plasmodium* infection (Raulf et al., 2019). Thus, Clec12a represents a promiscuous CLR with a variety of roles in immunity and unique signaling capabilities, yet its role in the intestine remains unknown.

We have previously demonstrated that specific gut fungi exacerbate colitis by inducing the purine metabolite, uric acid (Chiaro et al., 2017). Moreover, purine metabolism was recently associated with human intestinal disease (Zhu et al., 2019). These data collectively suggest that uric acid detection may be relevant to IBD. Based on this, we set out to explore the function of Clec12a in the intestine. We found that Clec12a deficient animals are highly susceptible to colitis that is dependent on the microbiota composition formed in the absence of Clec12a. FMT of Clec12a^-/-^ microbiota into WT germfree animals was sufficient to exacerbate colitis and formation of the colitogenic microbiota in Clec12a^-/-^ animals is independent of adaptive immunity. To identify relevant microbes that might exacerbate colitis, Clec12a^-/-^ animals were treated with various antibiotics and those that targeted gram-positive bacteria ameliorated disease. 16S rDNA sequencing identified *Faecalibaculum rodentium* to be significantly more abundant in Clec12a^-/-^ animals and oral gavage of *F. rodentium* was sufficient to exacerbate disease pathology in WT animals. Immune profiling of Clec12a^-/-^ animals revealed a marked reduction in homeostatic myeloid cells in the colon. RNA sequencing of *F. rodentium* treated macrophages revealed that Clec12a promotes phagocytosis. As Clec12a has been shown to bind multiple ligands, we asked if Clec12a would bind to diverse members of the microbiota. Using a Clec12a-Fc fusion protein we identify that Clec12a binds most highly to gram-positive members of the microbiota. Collectively these data suggest a model whereby Clec12a controls colitogenic members of the microbiota through direct binding and phagocytosis while preventing inflammatory responses. In scenarios where Clec12a expression is reduced, gram-positive pathobionts can expand and promote inflammation. Thus, our studies identify Clec12a as an important inflammatory suppressor to ensure homeostasis and maintenance of a healthy microbiota.

## RESULTS

### Clec12a^-/-^ animals are highly susceptible to colitis independent of the adaptive immune system

Reduced Clec12a expression has been reported in individuals with IBD and we have demonstrated that blocking the production of one of the ligands for Clec12a, uric acid, can ameliorate colitis (Begun *et al*., 2015; Chiaro *et al*., 2017; Liu *et al*., 2015). However, Clec12a has not yet been studied in animal models of colitis. To this end, WT and Clec12a^-/-^ C57BL/6 mice were challenged with acute DSS colitis. Clec12a^-/-^ animals lost significantly more weight, exhibited shorter colons and had a significant increase of the inflammatory marker LCN2 in feces when compared to WT animals, indicating worsened inflammation and disease (Figure 1A-C). Supporting this, histological analysis revealed increased epithelial damage, crypt loss, and influx of immune cells in Clec12a^-/-^ animals compared to WT (Figure 1D, E). Consistent with the results in the DSS colitis model, transfer of CD4^+^CD45RB^hi^ cells into TCRβ^-/-^;Clec12a^-/-^ double knockout mice led to less weight gain over time and shorter colons when compared to T cells transferred into TCRβ^-/-^ mice (Figure 1F, G). Collectively, these results demonstrate that loss of Clec12a leads to worsened intestinal disease.

**Figure 1.**
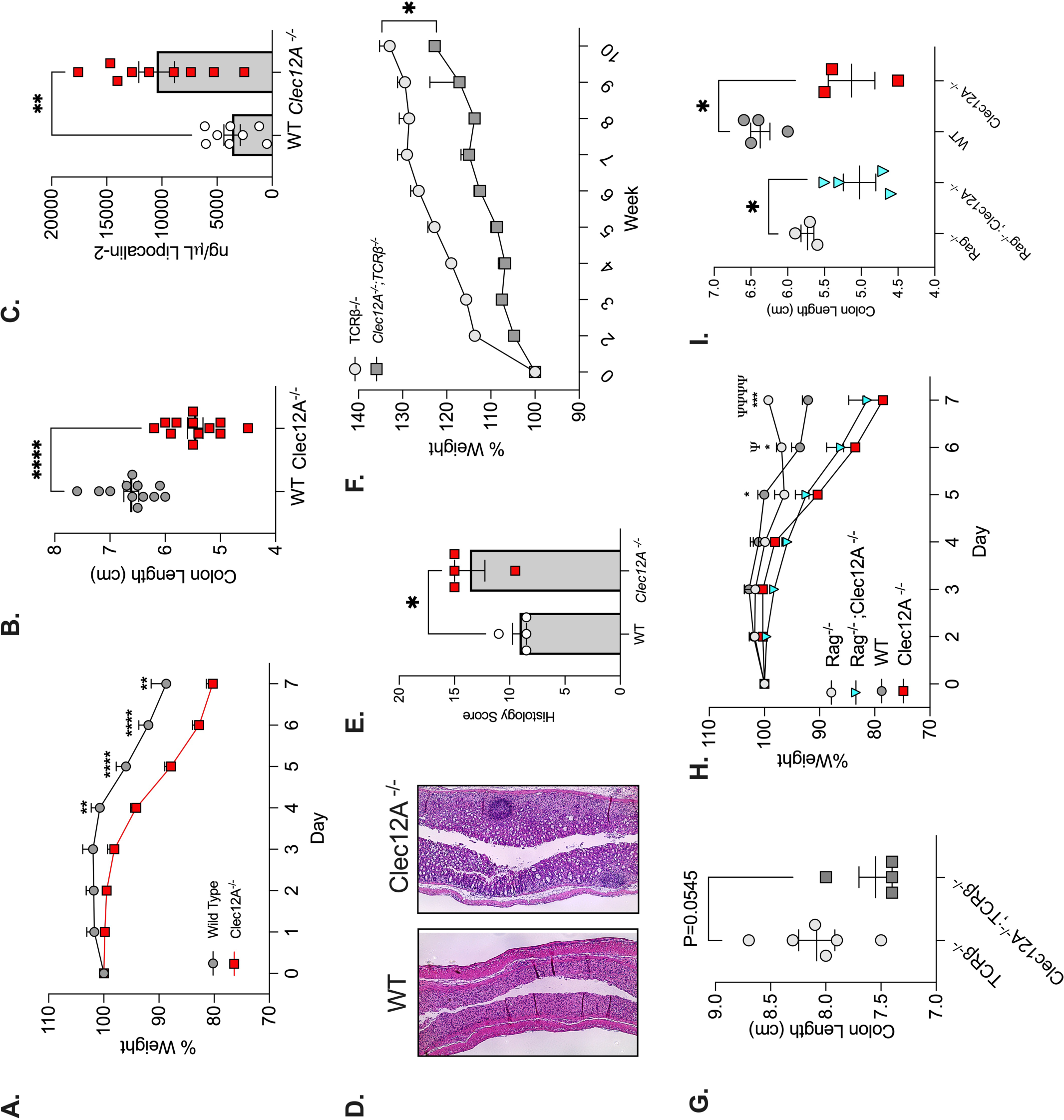
Clec12a restrains intestinal disease independent of the adaptive immune system. (A) In order assess colitis severity WT or Clec12a^-/-^ C57BL/6 specific pathogen-free (SPF) mice were treated with 2.5% dextran sodium sulfate (DSS) ad libitum for 7 days in the drinking water. Percent weight loss was assessed daily. Mean +/− SEM, data are pooled from three independent experiments using Two-way ANOVA and Śídák’s multiple comparison test. (B) Colon length (cm) of indicated animals was measured at sacrifice. Scatter plot is the mean +/− SEM, data are pooled from three independent experiments using unpaired t test. Each point represents one mouse (C) Fecal Lipocalin-2 (ng/mL) of indicated animals was measured as a readout of inflammation. The bar graph is the mean +/− SEM, data are pooled from two independent experiments using unpaired t test. Each point represents one mouse (D) Representative H&E stained colon sections from indicated animals after DSS-induced colitis. (E) Histology scores for indicated animals were generated from colon sections in (D). The bar graph is the mean +/− SEM, compared using unpaired t test. Each point represents one mouse. (F) T cell transfer model of colitis was performed to assess chronic colitis. Weight of TCRb^-/-^ and Clec12a^-/-^;TCRb^-/-^ animals was measured weekly. Graph depicts percent weight loss of indicated animals throughout the experiment. Mean +/− SEM, compared with 2-way ANOVA. (G) Colon length of indicated animals from (I) was measured at sacrifice. Bar graph is the mean +/− SEM, compared using unpaired t test. Each point represents one mouse. (H) WT, Clec12a^-/-^, Rag^-/-^ or Rag^-/-^ ;Clec12a^-/-^ C57BL/6 SPF mice were assessed for acute colitis. Indicated animals were provided with 2.5% DSS ad libitum for 7 days. Percent weight loss was assessed daily. Mean +/− SEM, data are representative of two independent experiments. Two-way ANOVA and Śídák’s multiple comparison test. (I) Colon length (cm) of indicated animals was measured at sacrifice. The bar graph is the mean +/− SEM, data are representative of two independent experiments using unpaired t test. Each point represents one mouse * p < 0.05; ** p < 0.01; *** p < 0.001; **** p < 0.0001

Clec12a is reported to largely be expressed within innate immune cells with very low expression reported in CD8+ T cells. Therefore, to determine whether adaptive immunity is involved in the worsened disease observed in Clec12a^-/-^ animals, Clec12a^-/-^ animals were crossed to Rag^-/-^ animals to remove both T and B cells. Again, Clec12a^-/-^ animals developed worsened disease when compared to WT animals as they lost more weight and had shorter colons than WT controls (Figure 1H, I). Similarly, Clec12a^-/-^; Rag^-/-^ developed worsened intestinal disease when compared to Rag^-/-^ controls and lost as much weight and had shorter colons than WT and Rag^-/-^ as Clec12a^-/-^ (Figure 1H, I). We have previously reported that oral treatment with *S. cerevisiae* enhances uric acid levels to worsen colitis. However, no differences were observed in uric acid levels of the serum or feces in WT versus Clec12a^-/-^ animals, suggesting that enhanced disease observed in the Clec12A^-/-^ is not a result of increased uric acid (Fig. S1). Thus, worsened colitis development observed in Clec12a^-/-^ animals is independent of the adaptive immune system.

### Worsened colitis in Clec12a^-/-^ animals is dependent on the microbiota

Colitis is known to be driven, in part, by changes to the composition of the microbiota (Kubinak et al., 2015; Ray and Dittel, 2015). To determine whether Clec12a^-/-^ animals harbor a microbiota that promotes worsened colitis, FMTs were performed from WT or Clec12a^-/-^ mice into germfree WT animals. Four weeks post-transplant, half of the animals were co-housed together, while the other half remained housed separately. Acute DSS colitis was then induced in each group (Figure 2A). Animals that received a Clec12a^-/-^ FMT and co-housed animals lost significantly more weight (Figure 2B) and had significantly shorter colons (Figure 2C) than animals that received a FMT from WT mice. Furthermore, histological analysis revealed greater epithelial damage and inflammatory influx after transplantation with a Clec12a^-/-^ microbiota (Figure 2D, E). The established microbial communities were distinct amongst the groups as determined by 16S rRNA gene sequencing using Bray-Curtis (PERMANOVA, p = 0.0001), unweighted (PERMANOVA, p = 0.0062) and weighted-Unifrac distances (PERMANOVA, p = 0.0001). The within group microbiota dissimilarity was greatest in co-housed animals by the same metrics indicating acquisition of microbes from each FMT genotype (Figure 2 F-H). These data indicate that worsened colitis within Clec12a^-/-^ animals is driven by specific microbes enriched in the absence of Clec12a; and these microbes became the dominate drivers of a colitogenic phenotype, even in the presence of an established WT microbiota.

**Figure 2.**
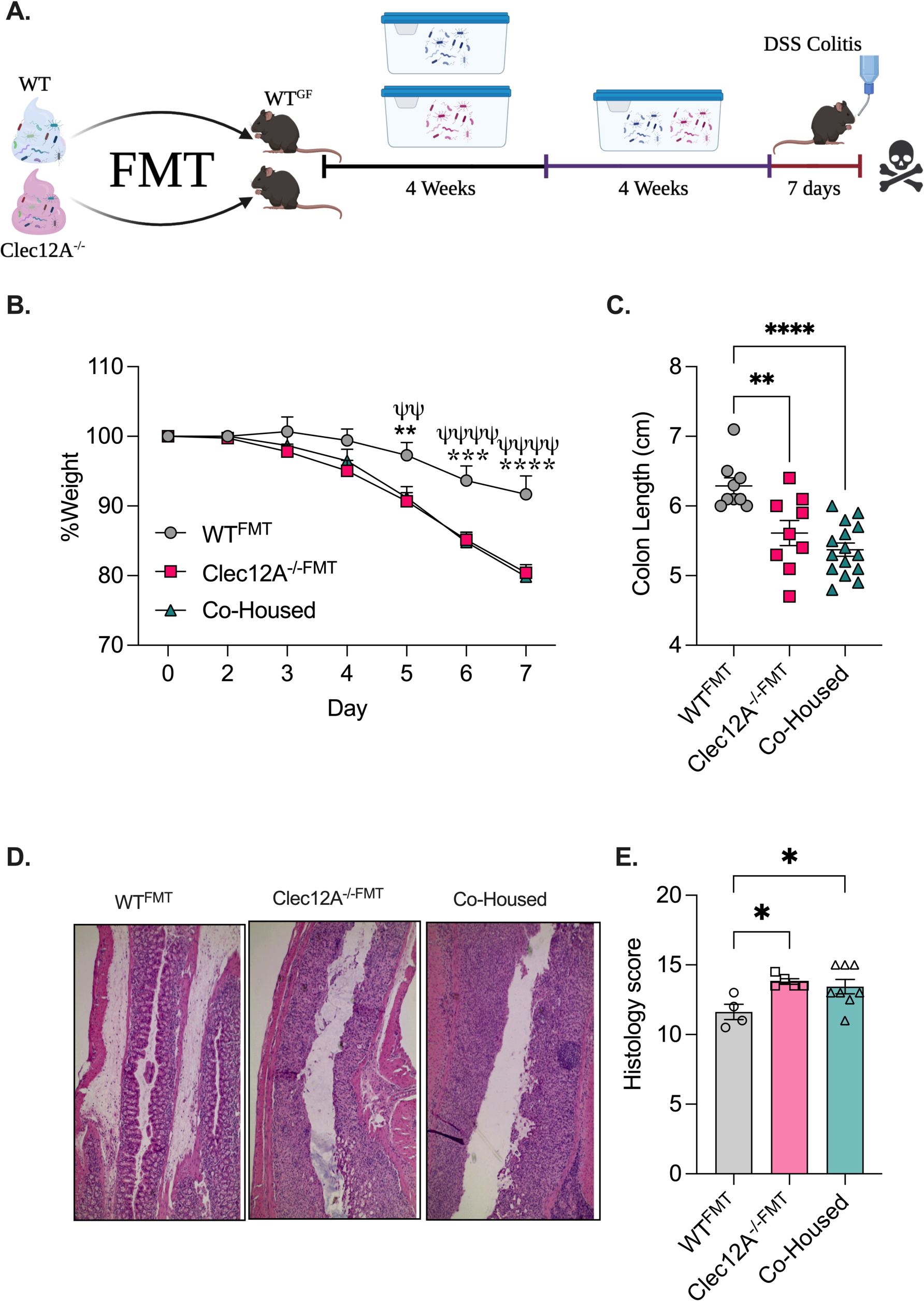
The microbiota formed in a Clec12a^-/-^ animals confers worsened colitis. **A)** Fecal microbiota transplants (FMT) were performed by transferring fecal supernatants from WT or Clec12a animals into germfree WT mice. Microbiotas were allowed to homogenize for four weeks, after which, half of the WT^FMT^ mice and half of the Clec12a^-/-FMT^ mice were co-housed and allowed another four weeks to mix. All groups then received 2.5% DSS ad libitum in their drinking water and were sacrificed on day 7. **B)** Percent weight loss of indicated animals was assessed daily. Graph depicts mean +/− SEM, data are pooled from two independent experiments using Two-way ANOVA and Śídák’s multiple comparison test. **C)** Colon length (cm) from indicated animals was measured at sacrifice. Scatter plot is the mean +/− SEM, compared using Ordinary One-Way ANOVA with Dunnett multiple comparisons test. Each point represents one mouse. **D)** Representative H&E stained colon sections of indicated animals after FMT and DSS-induced colitis. **E)** Histology scores were generated from colon sections in (D). Bar graph is the mean +/− SEM, compared using unpaired t test. Each point represents one mouse. **F)** Weighted Unifrac distances from 16S rRNA gene sequencing of WT^FMT^, Clec12a^-/-FMT^ or co-housed animals. Box and whisker plots showing all points (pairwise distance comparison) and compared using a Kruskal-Wallis test. **G)** Unweighted Unifrac distances from 16S rRNA gene sequencing of WT^FMT^, Clec12a^-/-FMT^ or co-housed animals. Box and whisker plots showing all points (pairwise distance comparison) and compared using a Kruskal-Wallis test. **H)** Bray-Curtis distances from 16S rRNA gene sequencing of WT^FMT^, Clec12a^-/-FMT^ or co-housed animals. Box and whisker plots showing all points (pairwise distance comparison) and compared using a Kruskal-Wallis test.

### Microbiota composition formed in Clec12a^-/-^ animals is independent of adaptive immunity

The immune system is important for dictating the composition of the microbiota. The FMT experiment indicated that Clec12a^-/-^ leads to a change in the composition of the microbiota that promotes worsened colitis development. We first wanted to determine whether Clec12a^-/-^ deficiency within the immune compartment was capable of re-shaping the gut microbiota. To this end, bone marrow from WT or Clec12a^-/-^ mice was transferred into lethally irradiated WT SPF recipient mice in order to standardize the microbiota followed by an 8-week immune reconstitution period (Figure 3A). During this time, the transplanted immune systems have time to develop within the recipient, differentially shaping the composition of the microbiota. To determine whether pro-colitogenic changes to the microbiota resulted from this immune sculpting, we performed FMTs using the feces from either WT or Clec12a^-/-^ chimeric mice and induced DSS colitis (Figure 3A). While there were no significant changes in weight loss (Figure 3B), the colons from animals receiving a FMT from Clec12a^-/-^ chimeric mice were significantly shorter when compared to animals receiving the FMT from WT mice (Figure 3C), indicating worsened disease. Moreover, when WT and Clec12a^-/-^ FMT recipients were co-housed, the colons were significantly shorter when compared to animals receiving a WT FMT (Figure 3C). Thus, Clec12a^-/-^ deficiency within the hematopoietic compartment leads to the formation of a dominant colitogenic microbiota.

**Figure 3:**
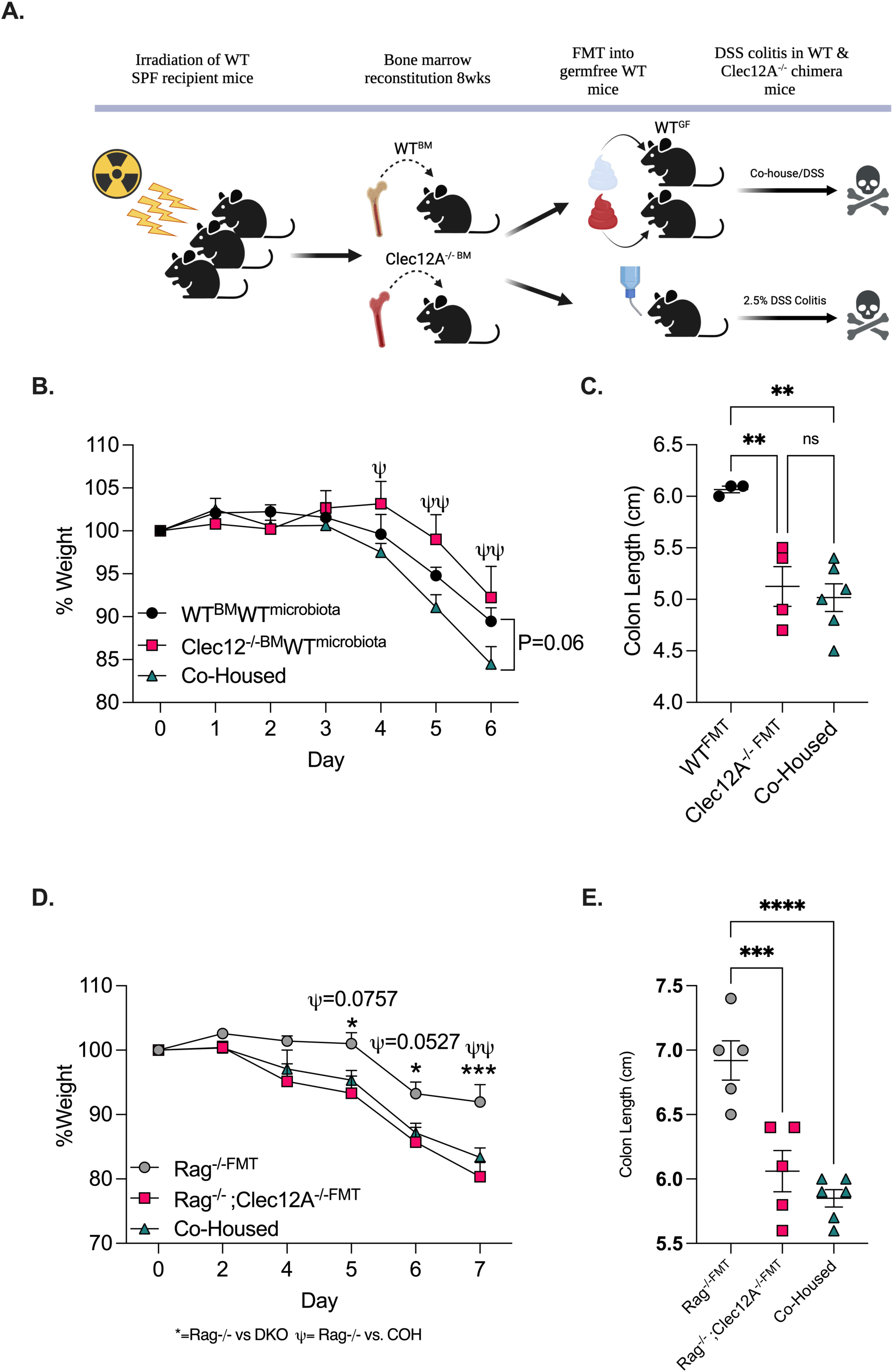
Microbiota composition formed in Clec12a^-/-^ animals is independent of adaptive immunity. **A)** Schematic diagram showing bone marrow reconstitution with subsequent FMT. C57Bl/6 WT mice were lethally irradiated and reconstituted with either WT or Clec12a^-/-^ bone marrow. After 8 weeks, feces were taken from these mice and transferred into WT germfree mice as explained in (A). The originally reconstituted chimeric animals were then provided 2.5% DSS ad libitum in their drinking water. **B)** Percent weight loss of FMT mice from (F) was assessed daily. Mean +/− SEM, using Two-way ANOVA and Tukey multiple comparison test. **C)** Colon length (cm) from indicated animals was measured at sacrifice. Scatter plot is the mean +/− SEM, unpaired t test. Each point represents one mouse **D)** FMTs to assess the role of the adaptive immune system on microbial shaping and colitis severity were performed by transferring fecal supernatants from Rag^-/-^ or Rag^-/-^;Clec12a^-/-^ SPF animals into WT germfree mice as described in (A). Mice were treated with 2.5% DSS for 7 days. Percent weight loss was assessed daily. Graph is the mean +/− SEM, using Two-way ANOVA and Tukey multiple comparison test. **E)** Colon length (cm) from indicated animals in (F) were measured at sacrifice. Mean +/-, SEM compared using Ordinary One-way ANOVA and Dunnet multiple comparison test. Each point represents one mouse. * p < 0.05; ** p < 0.01; *** p < 0.001; **** p < 0.0001

To further support that the adaptive immune system is dispensable for the formation of the colitogenic microbiota in Clec12a^-/-^ animals, we performed a FMT using feces from Rag^-/-^ or Rag^-/-^ ;Clec12a^-/--^ as described previously. Animals that received a Rag^-/-^;Clec12a^-/-^ FMT lost more weight and had significantly shorter colons than did animals who had received a FMT from Rag^-/-^ animals, moreover co-housed animals also developed worsened disease (Figure 3D, E). These data demonstrate that Clec12a deficiency within the innate immune system is sufficient to lead to the formation of a microbiota capable of aggravating colitis.

### Clec12a suppresses the expansion of *Faecalibaculum rodentium*

We utilized a series of different antibiotics to identify the relevant organisms that drive disease within the Clec12a^-/-^ animals. First, treatment with a potent cocktail of broad-spectrum antibiotics that include erythromycin, ampicillin, neomycin, and gentamycin, significantly reduced disease severity in both Clec12a^-/-^ and WT animals, suggesting that the bacterial population of the microbiota was relevant for disease (Figure 4A; Figure S2). Next, we further narrowed down which bacterial populations were most relevant for disease by utilizing different antibiotics targeting unique bacteria taxa. Specifically, we selected gentamycin, an aminoglycoside targeting gram negative bacteria, and cephalothin, a cephalosporin that primarily targets gram positive bacteria. These drugs were administered to Clec12a^-/-^ animals during DSS colitis. While treatment with gentamycin made no difference in disease severity, animals that received cephalothin maintained their weight and had significantly longer colons than mock treated animals (Figure 4B, C), indicating that gram positive organisms residing within the Clec12a^-/-^ microbiota play an important role in exacerbating disease pathology.

**Figure 4.**
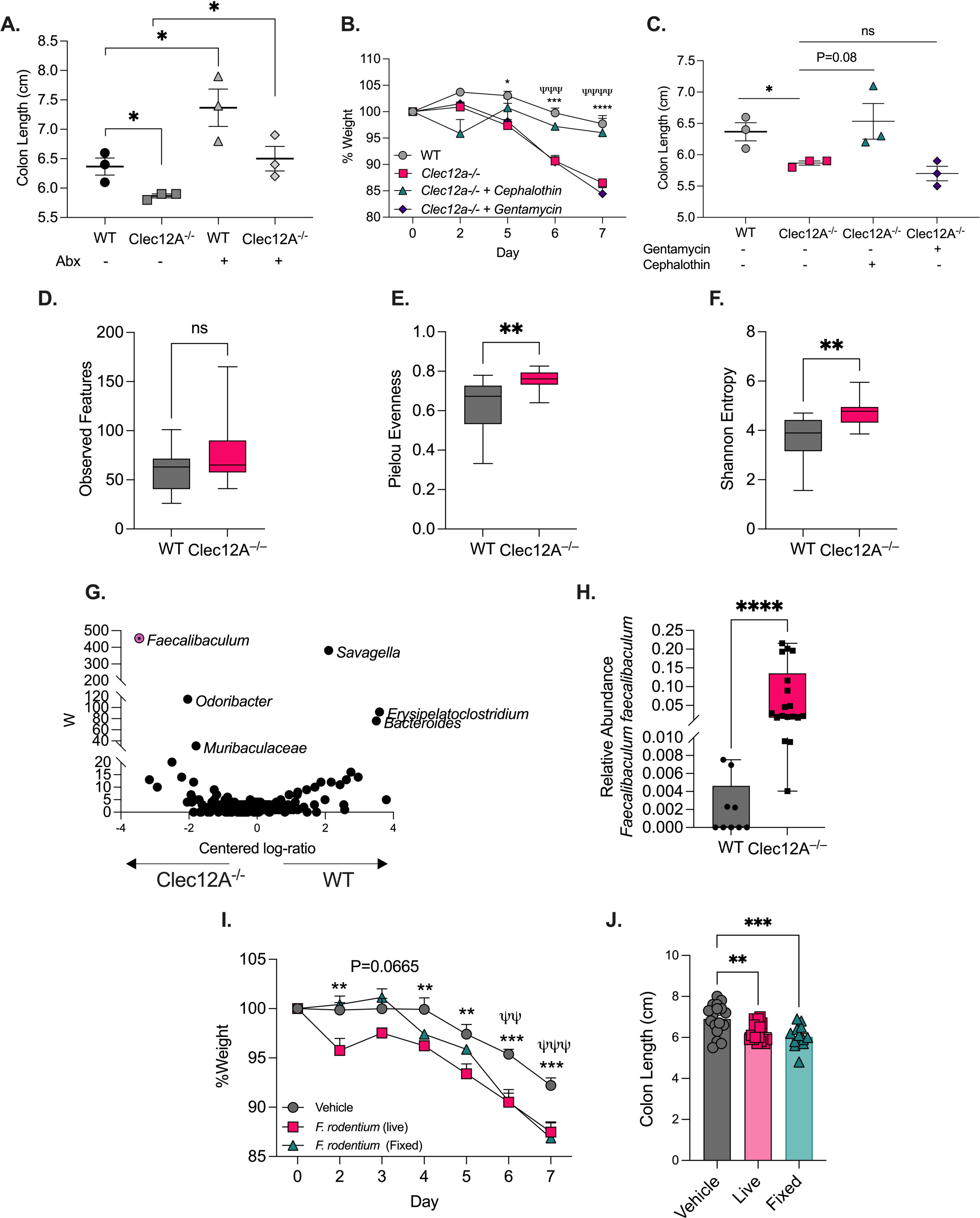
Clec12a limits the expansion of *Faecalibaculum rodentium,* a microbe that worsens colitis. A) WT or Clec12a^-/-^ C57BL/6 SPF mice were given an antibiotic cocktail (A) (neomycin, gentamycin, ampicillin, erythromycin), cephalothin, or gentamycin (B, C) at a concentration of 0.5g/L for fourteen days. After seven days animals were given 2.5% DSS in their drinking water until sacrifice. Colon length (cm) from indicated animals were measured at sacrifice. Scatter plot is the mean +/− SEM, unpaired t test. Each point represents one mouse. B) Percent weight loss of indicated animals was assessed daily. Graph is the mean +/− SEM, using Two-way ANOVA and Tukey multiple comparison test C) Colon length (cm) of indicated animals measured at sacrifice. Scatter plot is the mean +/− SEM, unpaired t test. Each point represents one mouse. D) Observed features from WT or Clec12a^-/-^ feces, 16S rRNA sequencing alpha diversity. Mean +/− SEM, compared using Mann-Whitney test. E) Pielou Evenness from WT or Clec12a-/- feces, 16S rRNA sequencing alpha diversity. Mean +/− SEM, compared using Mann-Whitney test. F) Shannon Entropy Mean +/− SEM, compared using Mann-Whitney test. G) ANCOM analysis from WT and Clec12a^-/-^ feces. Taxonomic names are best classifications of ASVs using the Genome Taxonomy Database (GTDB). H) Relative abundance of *Faecalibaculum rodentium* ASV’s. Compared using Mann-Whitney I) C57Bl/6 WT SPF mice were pre-treated with 1x10^8^ CFU of *F. rodentium* by oral gavage for three days. Animals then received 2.5% DSS in their drinking water for 7 days while continuing to received daily gavage of *F. rodentium.* Percent weight loss was assessed daily. Mean +/− SEM, Data are pooled of four separate experiments using Two-way ANOVA and Tukey multiple comparison test. J) Colon length (cm) of indicated animals was measured at sacrifice. Bar graph is the mean +/− SEM, Data are pooled of four separate experiments compared using Ordinary One-Way ANOVA with Dunnet multiple correction test. Each point represents one mouse. * p < 0.05; ** p < 0.01; *** p < 0.001; **** p < 0.0001

We next performed 16S rRNA gene sequencing of SPF WT and Clec12a^-/-^ feces to profile compositional changes in the microbiota. WT and Clec12a^-/-^ animals clustered significantly different by both non-phylogenetic and phylogenetic abundance-weighted metrics, Bray-Curtis and weighted-UniFrac, respectively (Figure S3A-C). Alpha diversity metrics revealed no significant differences in richness (observed features), indicating Clec12a^-/-^ did not harbor more species (Figure 4D). However, evenness and Shannon diversity indices were increased in the absence of Clec12a indicating that the proportional distribution of bacteria in Clec12a^-/-^ mice was altered (Figure 4E, F) and further supporting that Clec12a is important for regulating microbiota structure within the gut.

Analysis of composition of microbiomes (ANCOM) revealed that, among the ASVs (amplicon sequence variants) differentially abundant, an ASV classified in the *Faecalibaculum* genus (*Erysipelotrichaceae* family), was significantly expanded within the Clec12a^-/-^ feces compared to WT (Figure 4G, H). BLAST using the ASV representative sequence from our analysis indicated a 100% identity to sequences from the bacteria *Faecalibaculum rodentium*. These data provide evidence that gram-positive bacteria in the Clec12a^-/-^ animal are responsible for the colitogenic phenotype, and specifically implicate the gram-positive organism, *F. rodentium,* in worsened disease.

Based on this, we tested the hypothesis that *F. rodentium* might exacerbate colitis. We pre-treated WT SPF animals with 1 x 10^8^ CFU of either live or fixed *F. rodentium* by oral gavage and continued treatment every day during the acute model of DSS to ensure high levels of *F. rodentium* colonization. Animals that received both live and fixed *F. rodentium* lost more weight and had significantly shorter colons than vehicle treated animals (Figure 4I, J), indicating that increased levels of *F. rodentium* can worsen disease.

### Clec12a restricts macrophage inflammatory response while enhancing phagocytosis

To define Clec12a cellular expression in the absence of disease, we profiled innate and adaptive immune cells in the lamina propria (LP) using multi-color flow cytometry. Consistent with previous reports, Clec12a is predominantly restricted to the myeloid lineage with little to no expression on lymphocytes (Figure 5A) (Han et al., 2004; Pyz et al., 2008; Sobanov et al., 2001). Intestinal neutrophils, macrophages, and dendritic cells displayed similar levels of Clec12a expression (Fig. S4A), while macrophages had the greatest percentage of cells within the gut tissue expressing Clec12a (Figure 5A). Given the dual role of macrophages to promote inflammation via classical activation but also to initiate reparative mechanisms in response to tissue damage, we asked how Clec12a might influence this dichotomy during colitis. While macrophages are quite diverse, they can broadly be divided into M1 inflammatory and M2 reparative macrophages by expression of CD38 and EGR2 respectively. We show Clec12a^-/-^ animals have increased M1 macrophages and decreased M2 macrophages in the colonic LP during colitis (Figure 5 B, C, Fig. S4B). These data indicate that Clec12a regulates macrophage responses in the gut.

**Figure 5.**
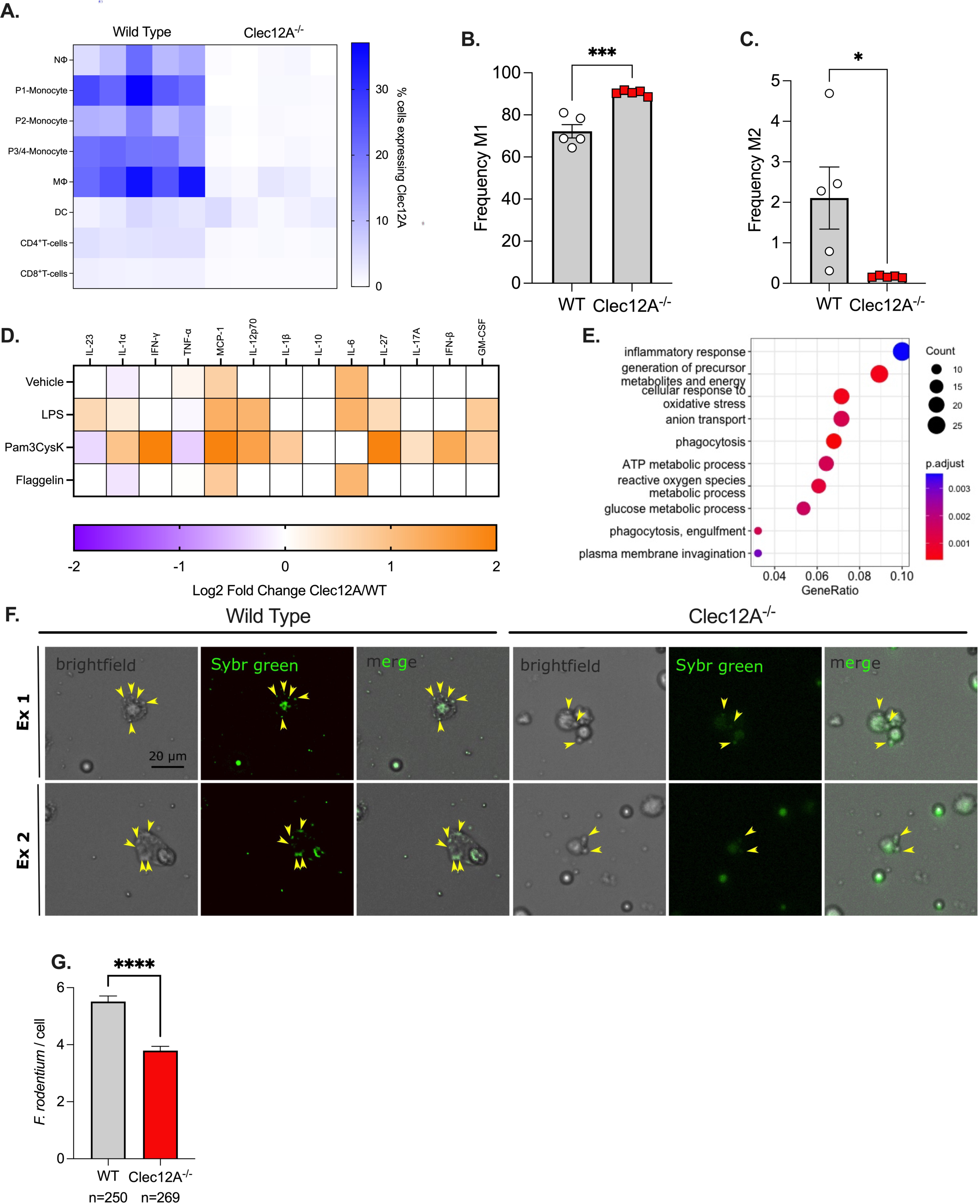
Clec12a limits inflammation and promotes phagocytosis in macrophages. A) Immunophenotyping was performed via flow cytometry on cells isolated from the colonic lamina propria (cLP) on WT and Clec12a^-/-^ animals. Heatmap shows frequency of Clec12a expression on innate and adaptive immune cells B) Bar graph shows the frequency of pre-gate, CD38^+^ cells (M1 macrophages) from the cLP of indicated animals. Mean +/− SEM, compared using unpaired t test. Each point represents one mouse. C) Bar graph shows the frequency of pre-gate, EGR2^+^ cells (M2 macrophages) from the cLP of indicated animals. Mean +/− SEM, compared using unpaired t test. Each point represents one mouse. D) BMDMs were harvested from WT and Clec12a^-/-^ long bones and cultured in the presence of M-CSF (20ng/mL) for 7 days. On day 8 BMDMs were pulsed with LPS (100ng/mL), MDP (10mg/mL), Pam3cysk (1mg/mL), or, Flagellin (1mg/mL) for 24hours. Media was collected and assayed for cytokine production. Heat map shows Log2 fold change of indicated cytokines from Clec12a^-/-^ over WT. E) BMDMs from WT and Clec12a-/- mice were isolated and cultured as in (A) and then co-cultured with *F. rodentium* at a M.O.I. of 10 for 24 hours. After BMDMs were washed, RNA was collected, and prepared for RNA sequencing. Top 10 simplified GO term (Biological process) categories enriched amongst Clec12a^-/-^ down-regulated F) Peritoneal macrophages were isolated from WT and Clec12a^-/-^ mice and then co-cultured with Sybr green-stained *F. rodentium* in order to assess phagocytosis. Left: WT macrophages. Right: Clec12a^-/-^ macrophages. Representative images from brightfield (left columns), fluorescence (middle columns), and merged (right columns) are depicted. Yellow arrowheads denote examples of *F. rodentium* cells engulfed by macrophages. scale bar = 20 μm. Images are representative of two separate experiments. G) Bar graph shows organisms per cell, seen in (F). 250 WT cells and 269 Clec12a^-/-^ cells were counted to quantify the amount of *F. rodentium* per cell.

To better understand how Clec12a influences macrophage biology in response to microbial ligands, bone marrow derived macrophages (BMDMs) isolated from WT or Clec12a^-/-^ animals were treated with a variety of common microbial ligands including LPS, Pam3CysK and Flagellin. Cytokine analysis was performed on all these samples. Many of the cytokines tested were not induced under these conditions, however MCP-1, IL-27, IFN-β, TNF-α and IL-6 were consistently detected in these cultures (Figure 5D). Clec12a^-/-^ BMDMs had significantly higher expression of these cytokines in response to many of the microbial ligands and the greatest differential responses were seen in response to the TLR2 agonist Pam3CysK. This data is consistent with the known role of Clec12a in tempering inflammatory responses during sterile inflammation (Neumann *et al*., 2014).

RNA-seq was performed on BMDMs from WT or Clec12a^-/-^ treated with *F. rodentium* to determine how Clec12a regulates responses of these cells to this specific bacterial organism. We assessed the differential response by comparing Clec12a^-/-^ BMDMs exposed to *F. rodentium* over vehicle controls to that of WT BMDMs. Enrichment analysis of differentially expressed genes showed that Clec12a deficient macrophages had a significant increase in genes involved within cell cycle control when challenged with *F. rodentium* (Figure S5A). Cell cycle is intrinsically linked with phagocytosis and there were also significant decreases in genes associated with phagocytosis and the inflammatory response (Figure 5E) (Luo et al., 2005). Consistent with a recent study on Clec12a and *Salmonella,* these data suggest that phagocytosis is defective in the absence of Clec12a in response to the normal gut resident *F. rodentium (Begun et al., 2015)*. To test this, peritoneal macrophages were collected from WT and Clec12a^-/-^ animals and co-cultured with Sybr®-green labeled *F. rodentium*. Clec12a^-/-^ macrophages had significantly fewer microbes per cell (Figure 5F, G; Figure S5B), indicating that Clec12a deficiency leads to uptake of fewer organisms and is consistent with reduced phagocytosis in the RNA-seq dataset. Collectively, these data show that Clec12a plays the dual role of mitigating overt inflammatory responses to common microbial ligands while promoting macrophage bacterial uptake.

### Clec12a binds preferentially to gram-positive commensals

As phagocytosis is initiated by binding of microbes followed by internalization and killing, we hypothesized that Clec12a was able to bind to specific members of the microbiota, such as *F. rodentium*. To address this possibility, a fusion protein containing the extracellular binding domain of murine Clec12a and the Fc portion of human IgG1 (Clec12a-Fc) was created (Figure 6A) (Maglinao et al., 2014; Neumann *et al*., 2014; Raulf *et al*., 2019). The protein was purified and screened for binding to a panel of diverse gram-positive and gram-negative bacteria that are known to be present within the homeostatic gut, including *F. rodentium*, *Roseburia spp., Bifidobacteria animalis*, *Akkermansia muciniphila*, *Bacteriodes uniformis*, and the fungal commensal *Candida albicans,* by flow cytometry. *Salmonella enterica* was also included as previous studies indicated that Clec12a could associate with *S. enterica* entry into the cell (Begun *et al*., 2015). Clec12a bound to varying degrees with several phylogenetically distinct bacteria, including *F. rodentium,* but not the fungi, *C. albicans* (Figure 6B). Clec12a bound most highly to the gram positive organisms *F. rodentium* and a member of the Lachnospiraceae family *Roseburia sp.*, and interacted the least with *B. animalis*, *B. uniformis* and *S. enterica* (Figure 6B), indicating the Clec12a can bind to specific members of the microbiota with a preference for gram-positive organisms. Taken together our data identify Clec12a as a mechanism employed by the immune system to survey the gut microbial community and prune the microbiota through direct binding and phagocytosis while preventing overt inflammatory responses to abundant gram-positive microbes which is critical to maintaining gut health.

**Figure 6:**
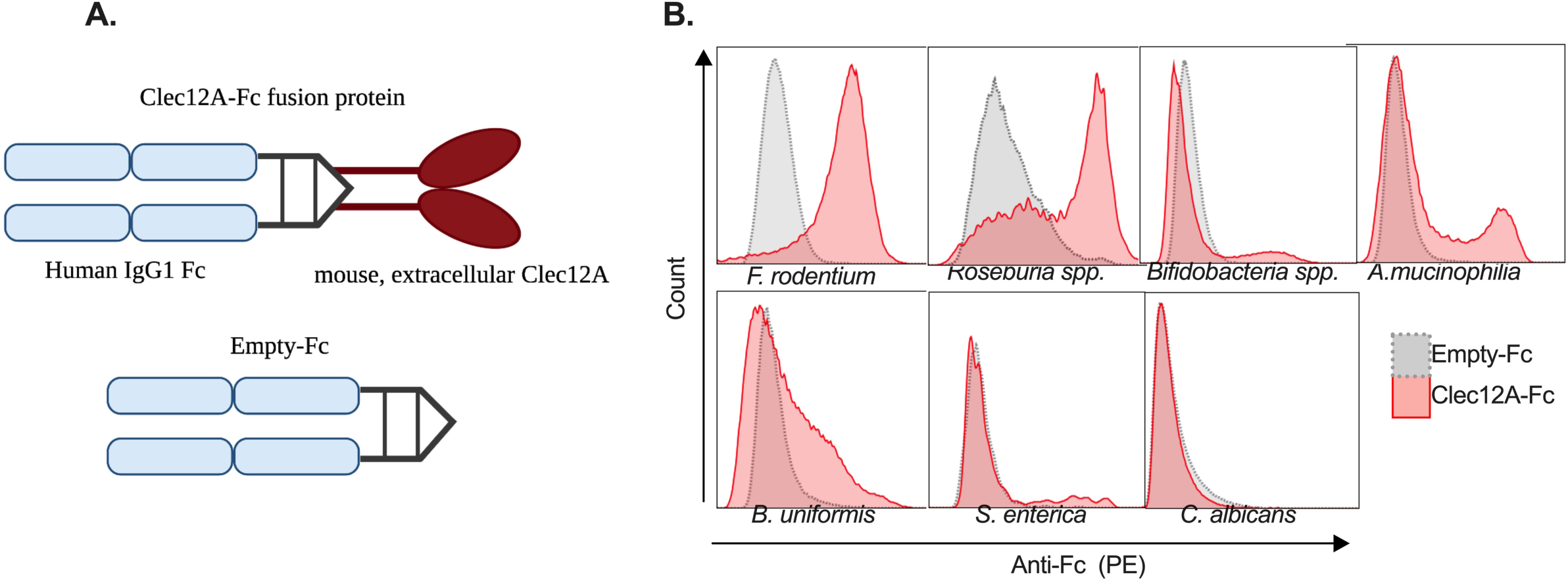
Clec12a directly binds to specific members of the microbiota. A) Schematic diagram depicting Clec12a-human IgG1-Fc fusion protein or empty vector (Fc portion only) B) Histograms of flow cytometric analysis representing differential binding of Clec12a-Fc to various bacteria or *C. albicans*.

## DISCUSSION

Homeostasis within the intestine requires mechanisms to restrict commensal microbes without initiating damaging inflammation (Macpherson et al., 2000). IgA, in recent years, has emerged as one of the critical ways in which gut immunity can exert control over the microbiota (Pabst and Slack, 2020). Indeed, IgA is able to directly bind to microbes and/or their products and eliminate them from the gut without the induction of proto-typical inflammation. Other immune molecules that possess these dual properties have not been studied to date. C-type lectins comprise a diverse class of proteins that can recognize a variety of protein, carbohydrate and lipid molecules from microbial and host products (Brown *et al*., 2018; Drouin et al., 2020). However, few have been studied in the context of gut immunity and the microbiota (Iliev *et al*., 2012; Martinez-Lopez *et al*., 2019). Thus, to our knowledge this is the first demonstration that Clec12a deficiency alters the composition of the microbiota that drives disease severity.

Clec12a has been largely been studied in the context of recognition of monosodium urate (MSU) or uric acid, which is often an indication of cellular death (Kono et al., 2010; Neumann *et al*., 2014). Uric acid is capable of activating inflammatory responses through the NLRP3 inflammasome, thus, in order to prevent tissue destruction during cellular death, this inflammation must be controlled (Martinon et al., 2006). Clec12a was shown to balance this inflammatory response though downregulation of Syk signaling (Marshall *et al*., 2004). Consistent with this, Clec12a deletion in a model of sterile inflammation and an autoinflammatory model of arthritis causes enhanced neutrophil activation and worsened disease, making Clec12a an important inflammatory brake (Redelinghuys et al., 2016). However, more recently Clec12a was also demonstrated to enhance type I interferon signaling in response to viral infection, and mediate uptake of *Salmonella* via the antibacterial autophagy pathway (Begun *et al*., 2015; Li *et al*., 2019). This demonstrates that Clec12a does not solely function to restrict inflammation, rather, can play a variety of roles depending on the context. The intestine is different in that there are a large number of immune cells even in the absence of disease. This is due to the presence of the microbiota, which plays an important role in instructing immune development and preventing colonization of pathogenic organisms. The microbiota possesses the same surface patterns detected by TLRs that also recognize pathogens, and therefore the intestinal immune system must have strict regulatory mechanisms in place to restrain inflammation to these foreign organisms (Blander et al., 2017).

We originally became interested in Clec12a because we identified that a gut fungi, *Saccharomyces cerevisiae* induces uric acid to worsen colitis (Chiaro *et al*., 2017). Moreover, other studies have identified that the microbiota may influence intestinal disease through induction of purine metabolism, of which the end product in mammals is uric acid (Zhu *et al*., 2019). These data suggest that mechanisms to sense uric acid, such as Clec12a, may influence intestinal homeostasis. However, we observed no changes in the levels of uric acid in serum or feces, indicating that enhanced disease severity in a Clec12a^-/-^ animal was not driven by increased uric acid. However, increased severity of colitis in Clec12a^-/-^ is driven by the composition of the microbiota. Thus, based on our data reduced expression of Clec12a^-/-^ could influence the composition of the microbiota in IBD patients (Jason L. Kubinak, 2015; Rehman et al., 2011; Vijay-Kumar et al., 2010). Interestingly, we found that antibiotics targeting gram positive organisms was able to ameliorate colitis severity in Clec12a^-/-^ animals. This is in contrast to several reports that have identified expanded gram negative Enterobacteriaceae during intestinal inflammation, thus, this is one of the few studies to have identified an expansion in gram positive organisms that is associated with colitis development. Although, likely not the only organism to influence colitis severity in this model, a dramatic expansion of *F. rodentium* was observed in Clec12a^-/-^ animals. *F. rodentium* is gram positive, oxidase and catalase-negative member of the Firmicutes (Bacillota) phyla (Chang et al., 2015). Introduction of *F. rodentium* daily to even WT mice enhances colitis, supporting that expanded levels of *F. rodentium* within the gut are harmful and likely driving colitis severity in the Clec12a^-/-^. A recent report demonstrated that *F. rodentium* can increase epithelial turnover and suppress expression of enzymes within epithelia that downregulate inflammatory responses and thus is in line with the worsened disease we see when *F. rodentium* is expanded in the gut (Cao et al., 2022). As humans are colonized with an organism similar to *F. rodentium* (Zagato et al., 2020), further studies should elucidate the influence of these *F. rodentium-*like bacteria on aspects of human physiology related to colitis.

Supporting a role for Clec12a amongst innate immune cells, we observed that the colitogenic microbiota is still formed in Clec12a^-/-^ animals that lack an adaptive immune system. The highest percentage of cells within the gut that express Clec12a were macrophages which led us to focus on Clec12a function in these cells. Previous studies have demonstrated Clec12a is an important mediator of type I interferon production in response to viral infection (Li *et al*., 2019). However, how Clec12a impacts responses to a broad spectrum of bacterial ligands had not been assessed; this is highly relevant in the intestine where an abundance of TLR ligands exist. Here, we show that Clec12a serves to temper macrophage reactions in the gut and likely functions to ensure homeostasis in response to the microbiota. RNA-seq of macrophages incubated with *F. rodentium* highlighted that Clec12a deficiency leads to cell cycle and phagocytosis defects in response to these commensal bacteria. A recent study showed that Clec12a^-/-^ macrophages had reduced autophagy in response to Salmonella (Begun *et al*., 2015). Our data compliments and extends these findings to include that Clec12a functions also to control non-invasive commensal organisms. *F. rodentium* has been shown by others to influence epithelial biology, which suggests that *F. rodentium* lies in close proximity to the host and thus must be tightly controlled by the immune system. Our data support that Clec12a controls *F. rodentium* though regulation of phagocytosis. Phagocytosis is initiated by physical binding of a ligand or microbe, and C-type lectins can associate with multiple ligands, which led us to test whether Clec12a can associate directly with gut-resident bacteria. Interestingly, Clec12a bound most highly to the gram-positive organisms *F. rodentium* and *Roseburia* and much less to the gram-negative organisms *A. muciniphila* and *B. uniformis* and not at all to *C. albicans.* This data shows that Clec12a has a propensity to bind specific members of the microbiota. This allows us to build a model whereby engagement of Clec12a and subsequent phagocytosis can control certain members of the microbiota and prevent the potentially detrimental effects of their expansion. This highlights the dual function of Clec12a that can restrain inflammation against microbial ligands while also recognizing and controlling specific members of the microbiota. Thus, Clec12a represents a unique way the innate immune system employs to ensure homeostasis within the gastrointestinal tract.

## METHODS

### Mouse strains

All mouse lines used herein were on a C57BL/6 background. Clec12a^-/-^ mice were kindly provided by Gordon Brown (Medical Research Council Centre for Medical Mycology, University of Exeter) and were genotype according to their previously described protocol (Redelinghuys, et al. 2016). *Rag2*^-/-^ and *TCRb*^-/-^ (Jackson Laboratories, 002118) were obtained from Jackson Laboratories and maintained in our mouse colony for several generations. For all *in vivo* colitis experiments 8 to 12-week old sex-matched mice were used. Both male and female mice were used. Mice were maintained in specific pathogen-free conditions.

### Bacterial and fungal strains

The following organisms were acquired from the DSMZ-German Collection of Microorganism and Cell Cultures GmbH: Faecalibaculum rodentium (DSM# 103405); Akkermansia mucinophilia DSMZ (DSM# 26127); Bifidobacterium animalis DSMZ (DSM# 26074). Bacteroides uniformis was acquired from the American Type Culture Collection (ATCC® 8492™). A Roseburia sp (strain designation JLR.KK002) was isolated on Schaedler’s agar from C57BL/6 mice harboring a spore former-enriched microbiota. Spore former enrichment was performed as previously described and mice were maintained for multiple generations with this microbiota in our gnotobiotic facility before JLR.KK002 was isolated from feces (Browne et al., 2016). We PCR-amplified and Sanger sequenced the 16S rRNA gene then classified it with the SILVA ACT service and SILVA database (Pruesse et al., 2012). Salmonella typhimurimum ST2 strain was used. Candida albicans strain sc5314 was used. Bacterial strains used in the PhyLo11b synthetic community are described below.

### DSS-induced experimental colitis

Colitis was induced with the addition of 2.5% or 3% (w/v) 36,000-50,000MW Dextran sodium sulfate (DSS) (MP Biomedicals) in the drinking water for 7 days. Colitis severity was monitored daily through changes in body weight, stool consistency, and fecal blood. For monocyte rescue experiment all mice received 2.5% DSS for 5 days, after which regular water was given; weight monitoring continued to day 10 when animals were harvested. Severity limits of greater than 20% weight loss were imposed as a severity threshold. At experimental endpoints, MLNs and colon were removed. The colon was measured from cecum to rectum to determine length. Tissues were then processed for histology, RNA extraction, or flow cytometric analysis.

### Colon Histology and Scoring

Colons were fixed within histology cassettes O/N at RT in 10% buffered formalin phosphate solution after removal of fecal contents and then transferred to 70% EtOH at 4 C until staining. Sectioning and H&E staining was done at the HCI Biorepository and Molecular Pathology Resource core in the ARUP operated Research Histology division. Colon scoring was done in a blinded manner with the following rubric scales: percentage of colon crypt loss (0.5 (5%) - 7 (>75%)), crypt loss severity (1 (partial loss) – 5 (full loss)) and inflammatory aggregates (1 (1-3) – 3 (>7)).

### Lipocalin-2 (LCN-2) ELISA

Protocol was performed according to manufacturer’s protocol (Fischer Scientific DY1857-05). Briefly, feces were collected, weighed, and mashed in HBSS 1X at a concentration of 100mg/mL. Samples were then spun at 50 x g for 5 min to remove large debris. The supernatant was then removed and placed in a new tube and spun at 8000 x g for 5 min in order to pellet bacteria. This supernatant was then diluted according to sickness of animals but generally ranged from 1:100-500 (healthy) – 1:500-1000 (DSS-treated).

### Uric acid measurement

The concentration was of uric acid was performed according to manufacturer’s instructions (Invitrogen™ Amplex™Red Uric/Uricase activity assay kit, cat# A22181). Briefly, blood was collected from WT or Clec12a^-/-^ mice by cardiac puncture into gold top serum separator tubes. Tubes were allowed to clot for a 30min and then spun at 1300 *x g* for ten minutes. Serum was removed and placed in 1.5ml Eppendorf tubes, flash frozen and stored at −80 degrees Celsius until testing. Fecal uric acid was performed by obtaining fresh fecal pellets from WT and Clec12^-/-^ mice. Pellets were weighed and recorded and then mashed in 1mL of 1x reaction buffer. After physical mashing, tubes were spun at 50 *x g* for 5 min to removed large debris. The remaining fecal slurry was assayed for uric acid.

### T-cell transfer experimental colitis

WT spleens were isolated and mashed in RPMI containing 10% fetal bovine serum, penicillin and streptomycin. Cells were spun at 367 x g to isolate cells. The resulting pellet was then suspended in 1x RBC lysis buffer for 5 min, in the dark, at room temperature. The resulting white blood cells were then passed through a Miltenyi CD4^+^ MACS column to isolate CD4^+^ T-cells. T-cells were then stained with CD4 and CDRB and sorted on a FACS Aria Fusion (BD biosciences). CD4^+^ CDRB^hi^ cells were collected and administered to TCRb^-/-^ or TCRb^-/-^;Clec12a^-/-^ mice intra-peritoneally (I.P.). Each mouse received 5 x 10^5^ cells. Colitis severity and development was monitored weekly by assessing weight loss. At the experimental endpoint, MLNs and colons were removed and colons were measured from cecum to rectum to assess disease severity.

### Isolation of leukocytes from the colonic lamina propria

Murine colons with attached ceca were harvested and placed in 1X PBS in 6-well plates on ice. Any visibly remaining fat and connective tissue was removed and the cecum clipped from colons. Colons were splayed open and mucus and feces were carefully removed from the colon before being placed back in the 6-well plate. Once all colons were splayed and scraped, each colon was diced on a petri dish lid using a razor blade and placed in a separate 50 mL tube. 10 mL of pre-warmed dissociation solution (1X HBSS w/o Ca+ and Mg+, 1.5 mM DTT, 10 mM HEPES and 30 mM EDTA) was added and the tubes briefly vortexed. Colons were incubated for approximately 25 min. at 37°C with shaking at 150 rpm until solutions were cloudy from IECs separation. Samples were shaken 3X and vortexed for 15 seconds then poured over a 100 mM filter on a 50 ml conical to remove non-immune cells such as IECs, and the filters rinsed briefly with 2 mL of ice cold 1X PBS. The remaining tissue was collected on the filter with forceps and move to a new 50 mL conical. 15 mL of pre-warmed digestion solution (1X HBSS with Ca+ and Mg, 5% FBS, 50 U/ml Dispase II [Millipore-Sigma, cat # 4942078001], 0.5 mg/ml DNAse I [Worthington Biochemical, cat # LS002139] and 0.5 mg/mL Collagenase D [Millipore-Sigma, cat # 11088866001]) was added, samples briefly vortexed, and subsequently incubated for 45 min. at 37°C with shaking at 150 rpm, or until solution became cloudy. Samples were then shaken 3X and vortexed for 15 seconds. To collect the remaining immune cLP cells, the digested tissue was then poured over a 40 mM filter on a 50 mL conical tube containing 10 mL 1X PBS. Samples are next rinsed and filtered with 2 mL of ice cold 1X PBS and spun at 800 x g for 10 min at 4°C. Next, the supernatant was removed and discarded with vacuum. Cells were then resuspended in 500 ml of complete RPMI, and counted with Trypan blue exclusion staining.

### Flow cytometry

Leukocytes isolated from the cLP were stained for viability using zombie dye 510 (AmCyan) for 10min at RT in dark. Samples were then Fc Receptor blocked with CD-16/32 for 20 minutes at 4°C in dark. After spinning down at 367 x g for 5 minutes and decanted cells were stained for 30 minutes in the dark at 4°C. All antibodies were used at a dilution of 1:250, unless otherwise noted. Myleloid panel consisted of CD45-BV421, CD11b-PerCpCy5.5, MHCII-BV605, CD64-BV711, CX3CR1-BV785, Ly6C-FITC, CD38-PE-Cy7, Ly6G-CF-594, Clec12a-PE (1:500), EGR2-APC (intracellular, 1:50). T cells were distinguished by: Zombie dye 510 (AmCyan), CD45-BV421, CD3e-BV711, CD4-FITC, Foxp3-APC (intracellular 1:50), IL-10-PE (intracellular 1:50), IFN-g (BV605) (intracellular 1:50), RORgt-PE610 (intracellular 1:50), IL-17A-BV785 (intracellular 1:50). After staining, cells were washed 2 times for 5 minutes each with column buffer (1X HBSS w/o Ca^2+^, Mg^2+^ and HEPES, EDTA, FBS). Cells were fixed O/N with 2% PFA if no intracellular staining was needed and then washed in column buffer 2X before running on the flow cytometer. If intracellular staining was necessary, the FoxP3 / Transcription Factor Staining Buffer kit (Tonbo Biosciences) and protocol was followed. All samples were then resuspended in 300 μl of column buffer and ran on a BD LSR Fortessa after compensating with UltraComp eBeads (ThermoFisher Scientific) and using single stain controls.

### IgA staining bacteria

Bacteria were isolated from fecal pellets by mashing into HBSS w/ Ca^2+^ and Mg^2+,^ at a concentration of 100 mg feces per mL HBSS. Mashed feces were then spun at 50 x g for 5 min to remove large debris. Supernatant was removed and placed in a new tube. This was then spun at 8000 x g for 5 min to pellet bacteria. Supernatant was removed and pellets were washed in HBSS w/ Ca^2+^ and Mg^2+^. Finally, pellets were resuspended in HBSS at a concentration of 100mg/mL and 10uL of this was placed in 96-well plate and blocked with HBSS w/ Ca^2+^ and Mg^2+^ containing 10% FBS for 20min at RT. 100uL of anti-IgA (PE) was then added directly to the sample and incubated for 30min at 4 C in the dark. Cells were then washed in HBSS+FBS solution 2x and lastly resuspended in HBSS+1x Sybr®-green and allowed to incubate for 20min in the dark at RT. Samples were then directly read on the flow cytometer.

### Bone marrow isolation

Bone marrow was isolate as previously described. Briefly, long bones from C57Bl/6 Wild type or Clec12a-/- animals were extracted and cleaned with a paper towel by gently rubbing. Clean bones were kept on ice while animals were processed. The end of the bones were slightly clipped in order to expose the marrow. Bone were inserted (exposed end facing down) into a 0.5mL Eppendorf tube which had been pierced with a 18g needle. This 0.5mL Eppendorf tube was then place inside a 1.5mL Eppendorf tube and spun at 13,000 x *g* for 1min. Spun through bone marrow was then RBC-lysed, using RBC-lysis buffer (Biolegend) according to manufacturer recommendations. After RBC lysis, cells were resuspended in complete Dulbecco’s Modified Eagle Medium (DMEM) containing: 10% FBS, 1mg/mL Pen/Strep, HEPES 10mM, 1x non-essential amino acid, Sodium Pyruvate 1mM. Cells were then spun at 367 *x g* for five minute to pellet cells. Isolated cells were then used to develop BMDMs, or to further isolate monocytes.

### Bone marrow derived macrophages

Leukocytes isolated from bone marrow isolation, suspended in complete DMEM and counted via trypan blue exclusion staining on a hemocytometer. Cells were plated on tissue culture treated 10cm dishes at a concentration of 5-8 x 10^6^ cells/plate in 10mL of complete media + M-CSF (20ng/mL). Fresh media, including M-CSF was added to existing media on days 4 and days 7. Fully differentiated BMDMs were used for experimental use between days 7-9.

### Monocyte Isolation

Bone marrow was harvested from WT and Clec12a^-/-^ animals as previously described. Monocytes were isolated with the Monocyte Isolation Kit (BM), mouse (Miltenyi Biotec) and an LS MACS column according to manufacturer instructions.

### *F. rodentium* BMDM co-culture and RNAseq

*F. rodentium* cultures were quantified by O.D. 600nm (using 1 OD 600nm = 1 x 10^8 cells/mL), washed 2x in sterile PBS and co-cultured with 1 x 10^6^ cell/mL fully differentiated BMDMs at an MOI of 10 in 6-well tissue culture plates. After 24 hours of co-culture, media was collected and stored at −20 C until further use. BMDMs were washed with cold, sterile PBS, 2x. Qiazol (Qiagen cat# 79306) RNA-stabilization reagent, was then added directly to each well. Wells were scraped with a cell scraper and collected into 1.5 Eppendorf tubes and frozen at −20°C until RNA extraction. RNA extraction was performed using, Direct-zol RNA miniprep kit (Zymo Research cat# R2070) according to manufacturer instructions and sequencing libraries were prepared by the Huntsman Cancer Institute High-throughput Genomics core facility. Total RNA was first hybridized with the NEBNext rRNA Depletion kit v2 (NEB, cat# E7400) to diminish rRNA from the samples. Stranded RNA sequencing libraries were prepared using the NEBNext Ultra II Directional RNA Library Prep Kit for Illumina (NEB, cat# E7760L). Purified libraries were qualified on an Agilent Technologies 4150 TapeStation using a D1000 ScreenTape assay (Agilent, cat# 5067-5582 and 5067-5583). The molarity of adapter-modified molecules was defined by quantitative PCR using the Kapa Biosystems Kapa Library Quant Kit (Roche, cat#07960140001). Individual libraries were normalized to 5 nM in preparation for Illumina sequence analysis and sequenced on a NovaSeq 6000 with a paired-end 150 cycle sequencing run.

### Cytokine analysis

BMDM were isolated and grown as previously described. Fully differentiated BMDMs were plated at 1 x 10^5^ cells/well in 12-well tissue culture plates. BMDMs were stimulated with100ng/mL LPS; 1 ug/mL Pam3Cysk; or 1ug/mL flagellin. Cells were stimulated for 24 hours, media was collected and saved at −20 C for analysis. Cytokines were analyzed with LEGENDplex™ Mouse Inflammation Panel (13-plex) (Biolegend, cat# 740150). Data was analyzed using the LEGENDplex™ data analysis software.

### 16s rRNA gene sequencing

Fecal pellets or ileal contents were collected from individual mice and immediately frozen at −70°C in 2mL screw cap tubes containing 250 mg of 0.15 mm garnet beads (MoBio, cat# 13122-500). DNA was extracted using the Power Fecal DNA Isolation Kit (MoBio or QIAGEN), per kit instructions and included 2 cycles of 1 minute of bead beating at 4°C on a Mini-Bead-Beater 16 (BioSpec Products). The V3 and V4 regions of the 16S rRNA gene was amplified with a single round of PCR using primers that contained (described 3’ to 5’) the V3/4 region 16S rRNA gene targeting sequence, a 2-nucelotide pad followed by the Illumina primer sequences, an 8-nucleotide index sequence and the remaining Illumina adapter sequence. The V3/4 16S-targeting sequences were taken from Takahashi (Takahashi et al., 2014) and the Indices were taken from Kozich (Kozich et al., 2013). The full oligonucleotide sequences used were (indices denoted by Xs): Prok16SV34_For: AATGATACGGCGACCACCGAGATCTACACXXXXXXXXACACTCTTTCCCTACACGACGCTC TTCCGATCTTGCCTACGGGNBGCASCAG; Prok16SV34_Rev: CAAGCAGAAGACGGCATACGAGATXXXXXXXXGTGACTGGAGTTCAGACGTGTGCTCTTCCGATCTGCGACTACNVGGGTATCTAATCC. PCR cycling conditions were as follows: 98°C initial denaturation for 2 minutes; 26 cycles of 20 sec 98°C denaturation, 20 sec 51.5°C anneal, 20 sec 72°C extension; and a single final 72°C extension for 2 minutes. Each PCR (performed in triplicate for each sample) was done in a 25 μl volume using Q5 High-Fidelity 2X Master Mix (NEB, cat# M0492L), 5 pmol of each primer and 50 ng template DNA. After amplification, triplicate PCRs from each sample were pooled, 5 μl was run on an agarose gel to confirm amplification and the remaining volume was cleaned up using Axygen AxyPrep MAG PCR cleanup beads (Corning, cat# MAG-PCR-CL-50) diluted to 62.5% in water to efficiently remove any primer dimer that would be preferentially sequenced. Diluted beads were added at 1.8X volume of the PCR reactions, cleaned per manufacturer guidelines and the cleaned amplicons eluted with 25 μl 10 mM Tris-Cl, pH 8.0. Amplicons were then quantified with a picogreen dsDNA assay (ThermoFisher, cat #P11495) on a microplate reader, then the cleaned and indexed individual sequencing libraries were evenly multiplexed (by ng DNA) and sequenced on an Illumina MiSeq instrument in paired-end 300 cycle mode at the Huntsman Cancer Institute’s High-Throughput Genomics shared resource facility.

### Bioinformatics processing of sequence data

16S rRNA gene raw reads were demultiplexed allowing 0 mismatches in indices then processed and analyzed within the QIIME2 framework (Bolyen et al., 2019) as we have previously described (Stephens et al., 2021). In brief, demultiplexed and quality-filtered sequences were first trimmed of primer and linker sequences with the Cutadapt plugin, then denoised with DADA2 (Callahan et al., 2016; Martin, 2011). Taxonomies were then assigned to ASVs with the classify-sklearn method in the feature-classifier plugin, against the GTDB reference set (release89) trimmed to the amplified V3/4 region and trained with the fit-classifier-naive-bayes method (Bokulich et al., 2018; Parks et al., 2018; Pedregosa et al., 2011). Diversity metrics, distances and statistical significance of beta diversity groupings by permanova were calculated within QIIME2 (Anderson, 2001).

For RNA-seq of BMDMs, the raw reads were first quality and adapter-trimmed using trim galore then transcript counts were quantified using Salmon against the transcriptome from assembly GRCm38 (mm10) (Patro et al., 2017). Transcript level quantifications were read into R using the tximeta package and summarized to gene level counts before differential expression analysis with DESeq2 using the design formula ‘∼ Genotype + Treatment + Genotype:Treatment’ (Love et al., 2014; Love et al., 2020). Differential responses to *F. rodentium* represented the difference between each genotype’s response to live *F. rodentium* treatment over vehicle control treated cells. Significantly differentially expressed genes were defined as those with an adjusted p value < 0.05 and log2 fold-change > |0.58| (1.5 fold-change). Significantly down-regulated and up-regulated gene lists were then used to test for enrichment in gene ontology (GO) terms using the clusterProfiler package including the simplify function to reduce GO term redundancy and identify key biological processes (S, 2020; Wu et al., 2021).

### *F. rodentium* DSS challenge

*F. rodentium* was grown as previously described and quantified to 1 x 10^9^ CFU/mL in sterile PBS. *F. rodentium* was fixed by first washing 2x in sterile PBS and then resuspending in 1mL 10% Fomalin-PBS for 10 minutes. Cells were then washed two more times in 10mL sterile PBS to prepare for oral gavage. Non-fixed were washed like fixed cells and kept on ice until gavage. WT C57BL/6 SPF mice were orally gaged with 100uL of solution containing either PBS alone (vehicle), *F. rodentium* (live), or *F. rodentium* (fixed) from days −3 to 10. Animals were given 3.0% DSS on days 0-7; fresh DSS water was replaced every other day. Severity limits of greater than 20% weight loss were imposed as a severity threshold. At experimental endpoints, MLNs and colon were removed. The colon was measured from cecum to rectum to determine length. Tissues were then processed for histology, RNA extraction, or flow cytometric analysis.

### Phagocytosis assay

Peritoneal macrophages were isolated by injecting 5mL Sterile PBS into the peritoneal cavity. PBS was gently agitated while in the cavity every few second. After 2 min, the PBS was withdrawn using a 28 gauge needle and dispensed into cold complete DMEM. Cells were spun at 367 x g and washed in complete media. Cells were then counted with a hemocytometer and plated into 12-well tissue culture plates at a concentration of 2.5 x 10^5^ cells/mL. Cells were allowed to adhere overnight. The next day, wells were gently mixed by swirling and media was aspirated. Adherent cells were then washed twice with 500uL of cold, sterile, PBS. Pre-warmed complete DMEM containing *F. rodentium* at a M.O.I. of 10 was then added to adherent (macrophages). Plates were then gently spun at 200 x g, at room temperature for three minutes and then incubated under standard cell culture conditions (37 C, 5% CO2) for 90 min. *F. rodentium* was prepared as described for co-culture of BMDMs for RNA sequencing only, after washing, cells were resuspended in a 1x Sybr® green (Sigma-Aldrich CAS: 163795-75-3), at RT for 20 minutes. Bacterial cells were then then pelleted at 4000 rpm for 5 min and resuspended with complete DMEM and pre-warmed for peritoneal macrophages. After 90 min of co-culture, media was aspirated, and cells were washed twice with cold, sterile, PBS. Adherent cells containing Sybr® green stained *F. rodentium* were fixed with 2% paraformaldehyde (PFA) for 10 minutes. PFA was aspirated and sterile PBS was placed on cells. Cells were then imaged for phagocytosis of *F. rodentium*

### Macrophage Image Analysis

Brightfield and fluorescence images of *F. rodentium*-treated macrophage cultures were taken using an EVOS m7000 Imaging System (Thermo Fisher Scientific, Waltham, MA). Three to five separate randomized images were taken for each culture, which were carried out in triplicate. Images were blinded and quantified using the software Fiji (Schindelin et al., 2012). Individual bacteria and macrophages were easily distinguished based on size. The total number of bacteria that were associated with (touching or overlapping with) single macrophages was quantified for a randomized subset of cells within the image. Equal numbers of cells were quantified between genotypes (n>200).

### CLEC12A-Fc fusion protein construction and purification

The cDNAs encoding the extracellular domains of mouse Clec12a were amplified by PCR from white blood cells isolated from C57Bl/6 WT SPF bone marrow and confirmed by sequencing. The cDNAs were fused to the C terminus of human IgG1-Fc in the expression vector pFuse-hIgG1-Fc2 (InvivoGen). Constructs containing Clec12a or empty vector were transfected into Lx293 cells cultured with DMEM +10% FBS and Pen/strep (1mg/mL). Secreted fusion proteins were harvested from the media 24-48hr after transfection. Media was spun at 367 x g for 5 min to remove debris/dead cells, and the supernatant was frozen at −20 C until use. Clec12a-Fc or Fc only fusion proteins were isolated with Pierce™ Classic Magnetic IP/Co-IP Kit (Thermo Fisher Scientific, Catalog number: 88804), quantified with Pierce™ BCA Protein Assay Kit (Thermo Fisher Scientific Catalog number: 23227) and confirmed by western blot using anti-Clec12a antibody (Novus Biologicals NBP2-27346SS).

### Clec12a-Fc-binding assay

Purified constructs were diluted to 0.4 mg/mL in HBSS w/ Ca^2+^ and Mg^2+^. Bacteria were grown in their respective media and aerobic conditions and normalized by O.D. 600nm. Bacteria was washed with HBSS w/ Ca^2+^ and Mg^2+^ and blocked with HBSS w/ Ca^2+^ and Mg^2+^ +10% FBS for 20 min at RT. All staining procedures took place under atmospheric conditions. 1 x 10^7^ bacteria cells were placed in a round bottom 96-well plate, spun at 3,000 x g and decanted. 50uL of constructs was placed on bacteria cells, gently mixed and incubated at 4 C for 1 hour. Bacteria cells were washed with HBSS w/ Ca^2+^ and Mg^2+^ + 10% FBS, 2x. 50uL of goat anti-human Fc (PE) (ebioscience Catalog # 12-4998-82) was added to cells and incubated at 4 C for 1 hour, in the dark. Cells were then washed as before and placed in HBSS w/ Ca^2+^ and Mg^2+^ containing 1x Sybr® green (Sigma-Aldrich CAS: 163795-75-3). Cells were incubated at RT for 20 min in the dark and read immediately on a BD LSR Fortessa flow cytometer.

### Irradiation and bone marrow reconstitution

Bone marrow was harvested from WT and Clec12a-/- animals as previously described. Recipient mice received were lethally irradiated with 500 Rads on day −1, followed by 400 Rads on day 0. Four hours after last irradiation, recipient mice were placed under isoflurane anesthetization and 100uL of isolated bone marrow cells suspended in sterile PBS (1 x 10^8^ cell/mL) were delivered via a 28.5 gauge needle to the retro-orbital space. We were interested in the microbiota thus animals were not placed on antibiotics, and after 8 weeks fecal pellets were collected for microbiota transfer into GF animals.

## AUTHOR CONTRIBUTIONS

T.R.C. helped conceive the study, performed most of the mouse and immunology experiments, put the data together into figures and helped write the manuscript. R.B. set up germfree experiments and maintained all germfree animals. E.S.V helped with the immunology gut lamina propria experiments. K.S.O. helped with mouse harvests and provided insight and direction. K.M.B helped with mouse harvests and flow cytometry. M.C.N. helped with mouse harvests. W.V. provided help with building the construct for the fusion protein and purification. T.J. provided help with phagocytosis assays. J.H. analyzed phagocytosis microscopy slides. M.H. provided help with the purification and use of the fusion protein. K.A.K. provide bacterial isolates. R.O.C. provided immunology expertise and insight. W.Z.S helped conceive the study performed bioinformatics analyses, helped with figure preparation and statistics and helped write the manuscript. J.L.R. helped conceive and direct the study, oversaw experimental design and outcomes, organize collaborators and reagents, garner funding for the study and helped write the manuscript.

**Supplemental Figure 1.**
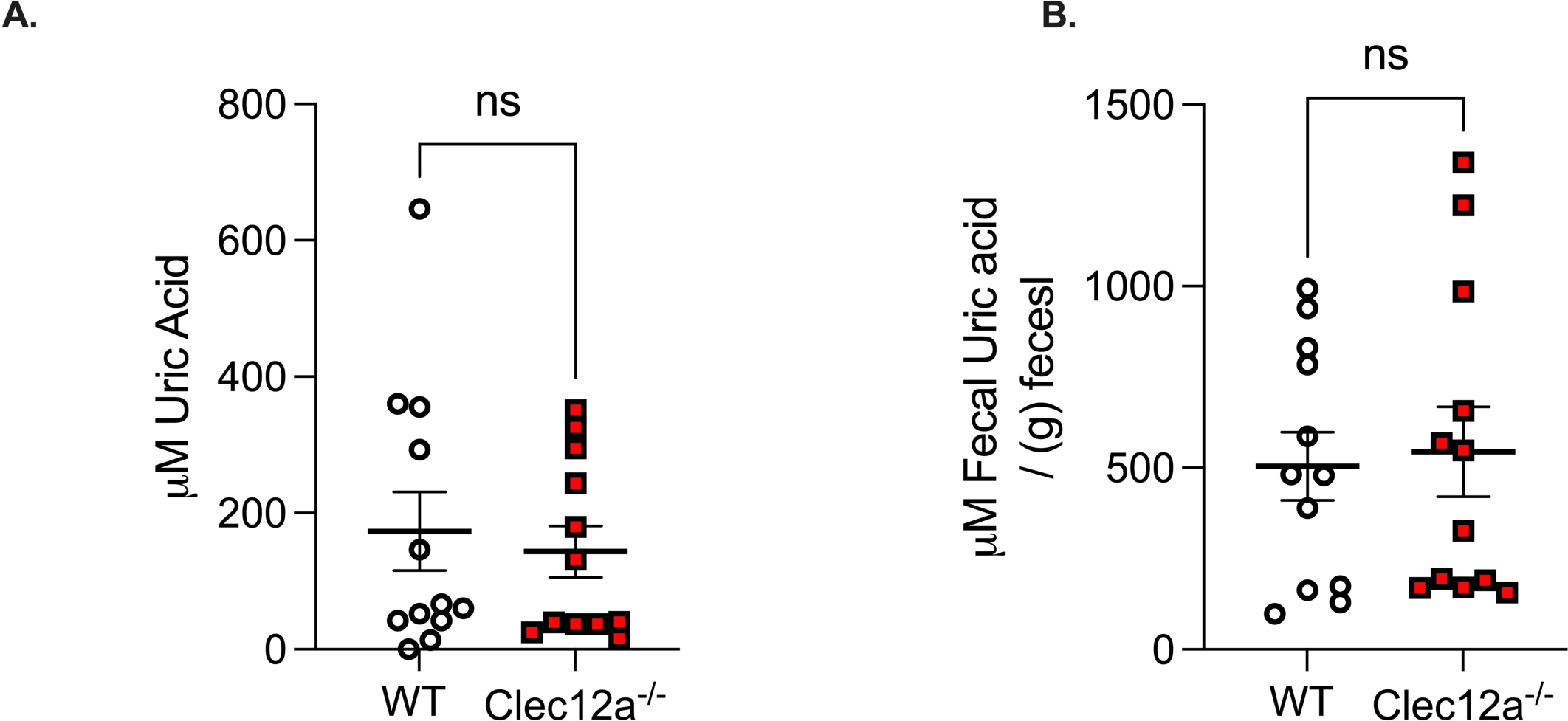
Absence of Clec12a does not disrupt uric acid levels. A) Serum uric acid (μM) of indicated animals was measured to assess purine metabolism between WT and Clec12a^-/-^ C57Bl/6 mice. The scatter plot is the mean +/− SEM, data are pooled from three independent experiments using unpaired t test. Each point represents one mouse. B) Fecal uric acid (μM) of indicated animals was measured to assess gut excretion of uric acid between WT and Clec12a^-/-^ C57Bl/6 mice. The scatter plot is the mean +/− SEM, data are pooled from three independent experiments using unpaired t test. Each point represents one mouse.

**Supplemental Figure 2.**
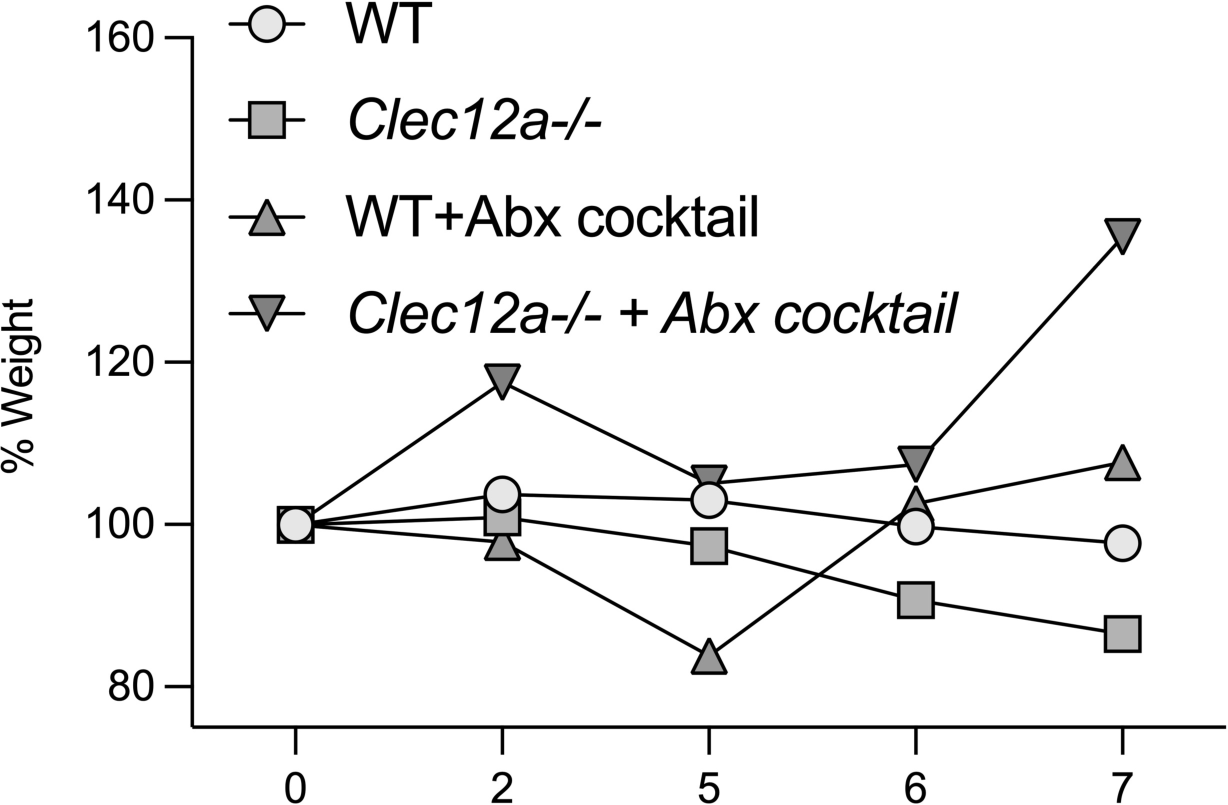
WT and Clec12a^-/-^ microbiotas drive differential disease. Percent weight loss of WT and Clec12a^-/-^ animals treated with antibiotic cocktail consisting of: neomycin, gentamycin, ampicillin, erythromycin.

**Supplemental Figure 3.**
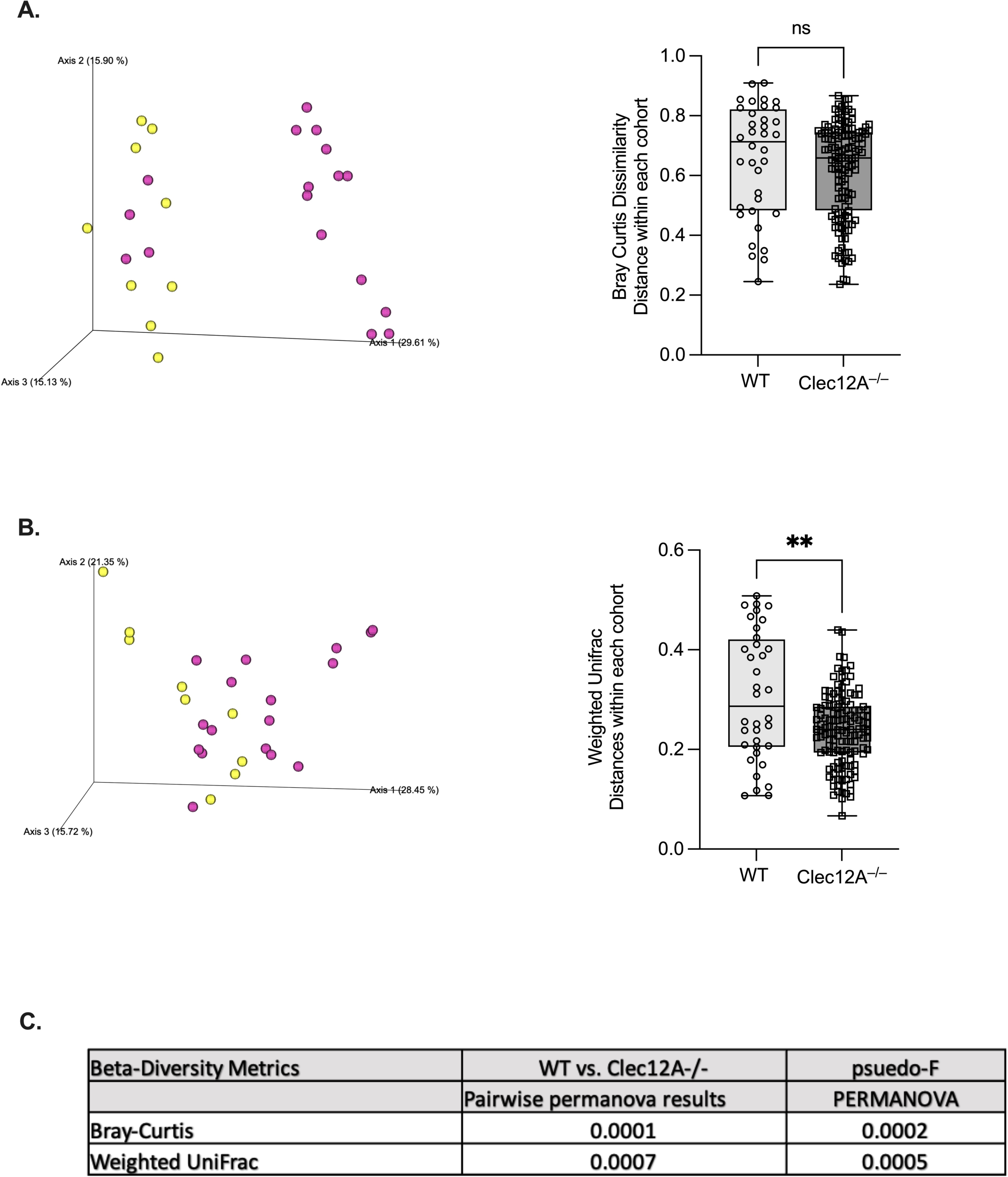
The microbiota composition is influenced by Clec12a. A) PCoA plots of first 3 axes based on Bray-Curtis distances showing overall microbiota relationships and the corresponding box and whisker plots quantifying dissimilarity within each group and compared with Mann-Whitney test. B) PCoA plots based on weighted Unifrac distances and the corresponding box and whisker plots quantifying within group dissimilarity compared using unpaired Mann-Whitney test. C) Table showing the PERMANOVA test results for WT vs. Clec12a^-/-^ fecal microbiota with Bray-Curtis and weighted Unifrac distances.

**Supplemental Figure 4.**
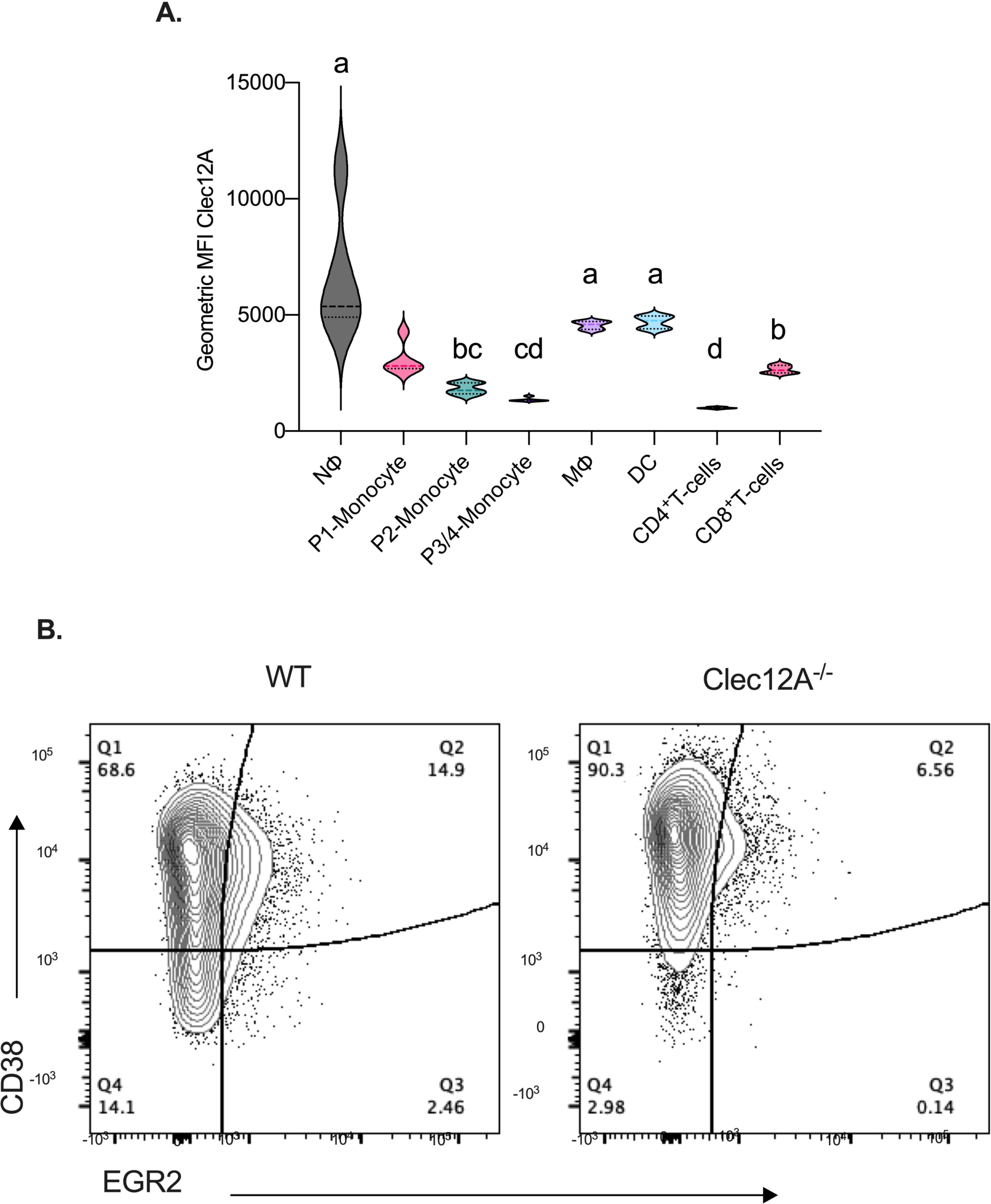
Clec12a in macrophages restricts inflammation. A) Violin plots depict the geometric mean fluorescent intensity (gMFI) of immune cells shown in (A). Mean +/− SEM, Kruskal-Wallis test with Dunn’s correction for multiple comparison. Letters denote groups that are significantly different. B) Representative flow cytometry plots of M1 and M2 macrophages from the cLP of WT or Clec12a^-/-^ mice after DSS colitis. Cells are pre-gated on live, CD45^+^, Ly6C^-^, CD11b^+^, MHCII^+^, CD64^+^. From Figure 5A

**Supplemental Figure 5.**
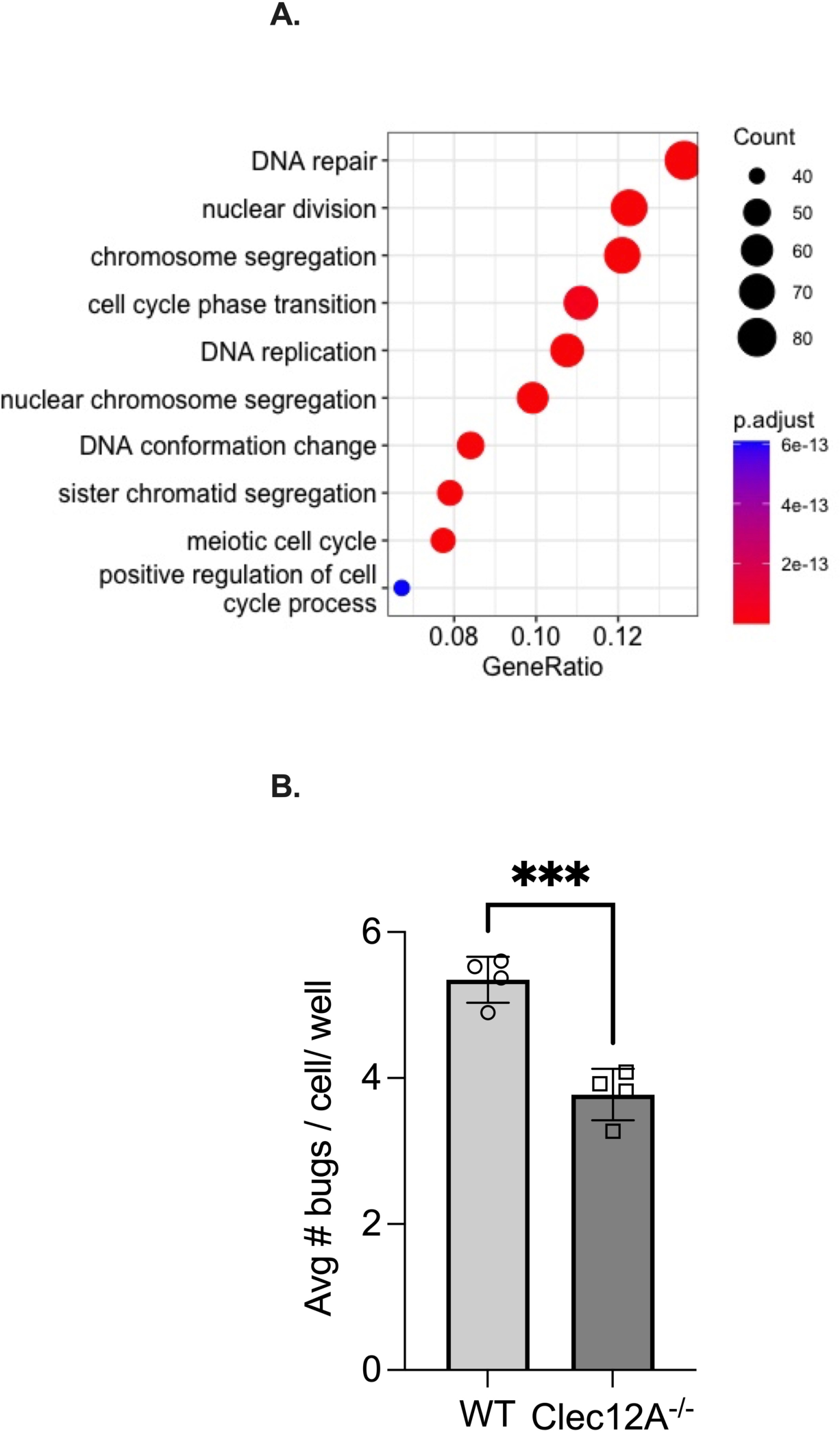
Clec12a stabilizes cell cycle control and regulates phagocytosis in macrophages. A) Top 10 simplified GO term (Biological process) categories enriched amongst Clec12a^-/-^ up-regulated genes compared to WT in response to *F. rodentium*. BMDMs from WT and Clec12a-/- mice co-cultured with *F. rodentium* were isolated and analyzed as in Figure 7A. B) Peritoneal macrophages were prepared as described and co-cultured with Sybr®-green stained *F. rodentium*. Bacteria cells were counted inside macrophages. >200 macrophages were counted. Bar graph shows the average number of *F. rodentium* from 4 individual wells to show consistency of counting and effect across wells. Mean +/− SEM compared using unpaired t test.

## ACKNOWLEDGMENTS

We would like to thank Gordon Brown for providing the Clec12a^-/-^ animals for this project. This work was supported by the Helen Hay Whitney Foundation (K.S.O), University of Utah NRSA Microbial Pathogenesis T32 Training Grant (K.S.O), a CCFA Senior Research Award (J.L.R), NIDDK R01DK124336, NIDDK R01124317 and NCCAM R01AT011423 (J.L.R), Edward Mallinckrodt Jr. Foundation (J.L.R), NSF CAREER award (IOS-1253278) (J.L.R), Packard Fellowship in Science and Engineering (J.L.R), Burroughs Welcome Investigator in Pathogenesis Award (J.L.R), American Asthma Foundation (J.L.R), Margolis Foundation (J.L.R), a MS Society Center grant (J.L.R), and the W.M Keck Foundation (J.L.R) NIH New Innovator Award DP2GM111099-01 (R.M.O), NHLBI R00HL102228-05 (R.M.O), an American Cancer Society Research Grant (R.M.O), a Kimmel Scholar Award (R.M.O), R01AG047956 (R.M.O), and NIAID R01AI141202 (A.S.I.). This work was supported by the University of Utah Flow Cytometry Facility in addition to the National Cancer Institute through Award Number 5P30CA042014-24. The support and resources from the Center for High Performance Computing at the University of Utah are gratefully acknowledged.

